# Temporal progression of anaerobic fungal communities in dairy calves from birth to maturity

**DOI:** 10.1101/2023.03.22.533786

**Authors:** Adrienne L. Jones, Jordan Clayburn, Elizabeth Pribil, Andrew P. Foote, Dagan Montogomery, Mostafa S. Elshahed, Noha H. Youssef

## Abstract

Establishment of microbial communities in neonatal calves is vital for their growth and overall health. Feed type and associated gastrointestinal tract morphophysiological changes occurring during the pre-weaning, weaning, and post-weaning phases are known to induce shifts in microbial community diversity, structure, and function. However, while the process has received considerable attention for bacteria, our knowledge on temporal progression of anaerobic gut fungi (AGF) in calves is lacking. Here, we examined AGF communities in fecal samples from six dairy cattle collected at 24 different time points during the pre-weaning (day 1-48), weaning (day 49-60), and post-weaning (3-12 months) stages. Quantitative PCR (qPCR) indicated that AGF colonize the calves GIT within 24 hours after birth, with AGF loads slowly increasing during pre-weaning/weaning phases and drastically increasing post-weaning. Culture-independent amplicon surveys identified higher AGF alpha diversity during pre-weaning/ weaning phases, compared to post-weaning. Further, the AGF community structure underwent a drastic shift post-weaning, from a community enriched in the genera *Khoyollomyces, Orpinomyces*, AL3, and NY8 (some of which commonly encountered in hindgut fermenters) to one enriched in the genera *Caecomyces, Piromyces, Pecoramyces, and Cyllamyces*, commonly encountered in adult ruminants. Inter-calf community variability was low in the pre-weaning/weaning phases but increased post-weaning. Finally, pairwise comparison of AGF community between calves day 1 post-birth and their mothers suggest a major role for maternal transmission, with additional input from cohabitating subjects. The distinct pattern of AGF progression in calves could best be understood in-light of their narrower niche preferences, metabolic specialization, and physiological optima when compared to bacteria, hence eliciting a unique response to changes in feeding pattern and associated structural development in the GIT of calves during maturation.

## Introduction

Beef and dairy cattle are key contributors to the global food supply. As herbivorous ruminants, cattle are largely dependent on their gastrointestinal tract (GIT) microbiome for nutrients acquisition. It is estimated that the cattle GIT microbiota provides 70% of their daily energy requirement via the fermentation of undigestible dietary substrates [1]. Cattle GIT is formed of the mouth, esophagus, a complex four-chambered stomach (reticulum, rumen, omasum, abomasum) leading to the small intestine and large intestine [2]. Microorganisms quickly colonize the GIT tract of cattle post-birth (see [3] for a recent review). In adult cattle, the rumen plays a key role in the digestive process. It acts as a fermentation chamber, where long resident time (50-60 hours [4]), anaerobic conditions, and stable temperature and pH enable the development of a complex resident community to effectively degrade ingested plant material to soluble products for absorption by the animal [2]. However, in newly born calves, the rumen is severely underdeveloped, while the omasum and abomasum represent the majority of the forestomach and are the sites of major digestive processes [5]. Calves receive milk or milk replacer diet during the pre-weaning phases, and these high protein substrates are mostly digested in the true stomach (abomasum) [5]. Weaning and gradual introduction of solid food induce a marked effect on GIT development, with the enlargement of the rumen at the expense of the omasum and abomasum. In addition to expansion, the absorption is stimulated by the development of papilla, which in turn is stimulated by the fermentation end product butyrate in a positive feedback loop [5].

Predictably, the progression of calves from birth to maturity and the associated nutritional shifts as well as GIT structural development lead to marked temporal shifts in the microbial community [5]. Understanding microbial community progression from birth to maturity in calves is of great interest from a basic, as well as and applied perspective, e.g., as a starting point for modulation of the microbial community towards higher productivity and better health outcomes. Patterns of temporal progression of the bacterial component of the calves GIT has received increasing attention [3, 6, 7], and a cumulative understanding of shifts in alpha and beta diversity patterns associated with calves weaning and maturation is emerging [3, 8–10]. However, in addition to bacteria and archaea, the GIT tract of cattle harbors a community of eukaryotes (fungi and protozoa). The anaerobic gut fungal (AGF) community in calves belongs to the phylum Neocallimastigomycota, and plays an integral role in nutrient acquisition, hence enhancing animal growth and overall health [11–17]. Our understanding of the scope of diversity and community structure of AGF in cattle and other herbivores is rapidly increasing [18–27].

However, to our knowledge, efforts to investigate the temporal progression of AGF load, diversity, and community structure in cattle from birth to adulthood have not been undertaken.

Here, we followed the progression of AGF community in six calves from birth to maturity. We quantified the AGF load using quantitative PCR (qPCR), studied their diversity, and community structure using culture-independent amplicon-based survey, and correlated the observed patterns to salient changes in calves’ feed and GIT development. We document clear, albeit unexpected, shifts in the AGF community associated with calves’ development that contrast patterns observed with the bacterial community. We argue that such differences are due to the unique niche requirements, specialized metabolic capacity, and narrow physiological preferences of AGF, compared to their bacterial counterparts.

## Materials and Methods

### Animal Management and Housing

All animal procedures were reviewed and approved by the Oklahoma State University Institutional Animal Care and Use Committee (Protocol #21-03). Calves (n = 6) were obtained from a commercial dairy (Braum’s Dairy Farm, Tuttle, OK, USA). All calves were born within approximately a 24-hour period ending the morning of May 6^th^, 2021. The farm personnel dipped the calves’ navels with iodine and provided colostrum prior to acquisition for the project. Calves were transported from the dairy facilities to the Ferguson Family Dairy Center at Oklahoma State University (Stillwater, OK, USA) on day 1 of the experiment. Upon arrival at the OSU dairy, the calves were ear-tagged and placed in pens (n = 3 per pen) with wheat straw bedding in a calf hutch. The straw bedding was replaced at least every 2 weeks while calves were housed in the hutches. On day 98 calves were moved to larger dirt pens but maintained in two groups of three calves. On day 180, calves were moved to a single pen with access to grass where they remained through the remainder of the experiment. At all times during the experiment, animals had ad libitum access to water.

### Feeding protocol

Calves were trained to drink milk replacer (Herd Maker 20-20 Bov Mos Medicated Milk Replacer, Land O’Lakes Animal Milk Solutions, Arden Hills, MN, USA) from a bucket feeder with nipples (Peach Teat; JDJ Solutions, LLC, Homer, NY, USA). Beginning on day 2, calves were fed at 07:30 and 16:00, receiving 1.89 liters of milk replacer (340 grams of milk replacer powder) per calf at each feeding. Beginning on day 4, each pen was provided with approximately 454 grams of textured calf starter feed (Ampli-Calf Starter 20; Purina Animal Nutrition; Arden Hills, MN, USA). Starter feed was monitored daily to ensure fresh feed was always available. On day 21, the milk replacer supplied was increased to 2.13 liters at each feeding. Calf starter feed was increased on day 27 to 680 grams per calf each day. On day 43 through day 48, milk replacer was reduced to 1 liter and was only fed once daily in the morning. calves were weaned from milk replacer at day 48. Additionally, on day 43, the starter feed supplied was increased to 454 grams per calf twice daily. Starter feed was increased to 1.36 kg/d for each animal on day 50 and then to 2.27 kg/d for each animal on day 60 with the daily allotment split between two feedings. Beginning on day 78, calves were given ad libitum access to Bermuda grass hay and starter feed was reduced to 1.13 kg/days for each calf. Calves were maintained on this feeding protocol until day 137, when the diet was switched to a grower ration supplied at 2.72 kg/day for each animal. The grower ration contained (percentage on dry matter basis): ground corn (43.32%), soybean meal (21.90%), oats (19.30%), alfalfa pellets (4.43%), molasses (4.29%), cottonseed hulls (3.92%), calcium carbonate (1.14%), trace mineral and vitamin mix (0.85%; included Rumensin 90 [Elanco Animal Health; Greenfield, IN, USA] to provide 0.14 grams/ton), and soybean oil (0.84%). Through the remainder of the experiment, calves were provided 2.72 kg/d for each animal of the grower ration and had ad libitum access to Bermuda grass hay. During the entire experiment, calves received vaccinations and occasional treatments as outlined in the supplementary document.

### Sampling

Fresh fecal samples originating from a single animal were obtained immediately post defecation and placed into sterile 50-ml plastic tubes and rapidly sealed. Samples were transferred to the laboratory on ice, usually within one hour, and stored at -20°C. Fecal samples were obtained from six calves at 24 different time points for a total of 144 samples. Samples were obtained at days 1 (within 18 hours of birth), 7, 14, 21, 28, 35, 42, 48 (pre-weaning phases), 49-55, 60 (weaning phases), and 120, 150, 180, 240, 270, 300, 330, and 360 (post-weaning phases). Dams were also sampled on the day of delivering calves (within 18 hours of birth) for dam-calf comparisons.

### DNA extraction

DNA extraction was conducted on a known weight of fecal material using DNeasy Plant Pro Kit (Qiagen®, Germantown, Maryland) according to manufacturer’s instructions. DNA from all samples was eluted in the same volume of TE buffer (50 μL).

### Quantitative PCR

Compared to unicellular bacteria and archaea, quantification of AGF in the herbivorous gut is problematic, given their complex life cycle (resulting in the coexistence of various stages at any given sampling time), wide variations in thallus morphology between genera (high biomass filamentous growth versus smaller bulbous thalli), drastic changes in AGF biomass pre- and post-feeding (with extremely low levels pre-feeding and a significant rapid increase post-feeding), and wide variability in DNA content/ biomass unit between polycentric and monocentric AGF taxa. Prior studies have quantified AGF load using spore counting [28], most probable number assay (thallus forming unit) [29], as well as protein content, chitin content, and chitin synthase activity in cell-free extract [30]. In addition, DNA-based qPCR methods have previously been used to quantify AGF using their SSU rRNA [31], their ITS1 region [25, 32–34], the more conserved 5.8S rRNA [35], and recently the D2 region of the LSU rRNA [36]. DNA-based qPCR methods have the advantage of higher sensitivity and circumventing the need for culturability.

qPCR reactions were conducted in 25-μl PCR reaction volume using the SYBR GreenER™ qPCR SuperMix for iCycler™ according to manufacturer’s instructions (ThermoFisher, Waltham, Massachusetts). The reaction contained one microliter of extracted DNA, 0.3 μM of primers AGF-LSU-EnvS primer pair (AGF-LSU-EnvS For: 5’- GCGTTTRRCACCASTGTTGTT-3’ and AGF-LSU-EnvS Rev: 5’-GTCAACATCCTAAGYGTAGGTA-3’) [36] targeting a ∼370 bp region of the LSU rRNA gene (corresponding to the D2 domain). Reactions were run on a MyiQ thermocycler (Bio-Rad Laboratories, Hercules, CA). The reactions were heated at 95°C for 8.5 min, followed by 40 cycles, with one cycle consisting of 15 sec at 95°C and 1 min at 55°C. A pCR 4-TOPO or pCR-XL-2-TOPO plasmid (ThermoFisher, Waltham, Massachusetts) with an insert spanning ITS1-5.8S rRNA-ITS2-D1/D2 region of 28S rRNA from a pure culture strain was used as a positive control, as well as to generate a standard curve. The efficiency of the amplification of standards (*E*) was calculated from the slope of the standard curve using the equation *E* =(10^-1/slope^) – 1, and was found to be 0.89. Number of copies of LSU rRNA genes in the samples were calculated from the standard curve and converted to copies/g feces based on the volume included in the reaction and the original weight of the fecal samples.

### PCR amplification, Illumina sequencing, sequence processing and phylogenetic assignments

PCR amplifications targeting the D2 region of the large ribosomal subunit (D2 LSU) utilized the AGF-specific primer pair AGF-LSU-EnvS primer pair [36], as described before [37]. The amplification protocol, amplicon cleanup, dual indices and Illumina sequencing adapter ligation, and library prep and pooling for Illumina sequencing were conducted as previously described [37]. Pooled samples were sequenced at the University of Oklahoma Clinical Genomics Facility using the MiSeq platform. Protocols for read assembly, and sequence quality trimming are described elsewhere [37]. Sequences were assigned to AGF genera as described previously [20, 37].

### Alpha diversity measures

Alpha diversity estimates (Shannon, Simpson, and Inverse Simpson diversity indices) were calculated using the command estimate_richness in the Phyloseq R package. ANOVA (calculated using the aov command in R) followed by post hoc Tukey HSD tests (using TukeyHSD command in R) was calculated to test the effect of day of sampling, weaning phase, and calf replicate in shaping the observed patterns of alpha diversity.

### Factors impacting AGF community structure

The phylogenetic similarity-based beta diversity index weighted Unifrac was calculated using the ordinate command in the Phyloseq R package. The obtained values were used to construct PCoA ordination plots using the function plot_ordination in the Phyloseq R package. PERMANOVA tests were run using the vegan command Adonis to partition the dissimilarity among the sources of variation (day of sampling, weaning phase, and calf replicate), and the F-statistics p-values were compared to identify the factors that significantly affect the AGF community structure. The percentage variance explained by each factor was calculated as the percentage of the sum of squares of each factor to the total sum of squares.

To assess the variability in community structure within each sampling day, or weaning phase, we first calculated group centroids using the vegan command betadisper. Following, the PCoA ordination distance of each sample to its group centroid was calculated (as the Euclidean distance between two points), and distances from group centroids were plotted in a box and whisker plot (using the command boxplot in R).

To identify genera differentially abundant in pre-weaning versus post-weaning phases, we calculated both linear discriminant analysis (LDA) effect size (LEfSe) [38], and Metastats [39] (in Mothur). Genera with calculated LDA scores and/or significant Metastats p-values were considered differentially abundant between the two stages.

### Sequence and data deposition

Illumina reads were deposited in GenBank under BioProject accession number PRJNA946728.

## Results

### Quantification of AGF load from birth to maturity

The presence of AGF was observed in all examined samples, including samples from calves at day 1 after birth. AGF load was extremely low at day 1 (6.78 x 10^2^ ± 6.02 x 10^2^ copies/g feces) (Figure 1A) and remained at comparatively low levels throughout the pre-weaning phase (day 1 – day 48) (2.47 x 10^3^ ± 2.43 x 10^3^ copies/g feces). A gradual increase in AGF load was observed during the weaning phase (day 49 – 60) (6.12 x 10^3^± 5.69 x 10^3^ copies/g feces) (Fig 1A-B). Post-weaning, a dramatic rise in AGF load was observed (Figure 1A-B), reaching 2.42 x 10^5^ ± 1.32 x 10^5^ copies/g feces at day 120, and remaining at comparable levels between day 150 and day 360 (3.02 x 10^5^ ± 2.31 x 10^5^ copies/g feces). Dams’ fecal samples taken on the day of delivery showed comparable AGF load to those observed in calves post-weaning (5.01 x 10^5^ ± 3.77 x 10^5^ copies/g feces) (Figure 1C). One-way ANOVA demonstrated that, while within weaning phases no statistical significance was observed between the different sampling days (Table S1), the weaning phase was found to significantly affect AGF load (F-statistic p-value = 1.7e^-19^, generalized eta squared (equivalent to the proportion of variability explained) = 0.49). Specifically, AGF load was significantly higher in the post-weaning phase when compared to the pre-weaning phase (Tukey post-hoc p-value: 5.28x10^-14^), as well as in the post-weaning phase when compared to the weaning phase (Tukey post-hoc p-value: 2.56 x10^-13^) (Figure 1b inset). On the other hand, the observed increase in AGF load between pre-weaning and weaning phases was not significant (Tukey post-hoc p-value=0.079) (Figure 1b inset). The same pattern held true when AGF load was compared between weaning phases within individual individual calves (Figure 1D, Table S2). On the other hand, AGF load did not differ significantly between the 6 calves within each weaning phase (Figure 1E). Collectively, these results demonstrate that AGF colonize the calves GIT within 24 hours of birth at very low thresholds, with their load slowly increasing during the pre-weaning and weaning phases, followed by a significant increase to numbers comparable to adult cattle post-weaning.

**Figure 1.**
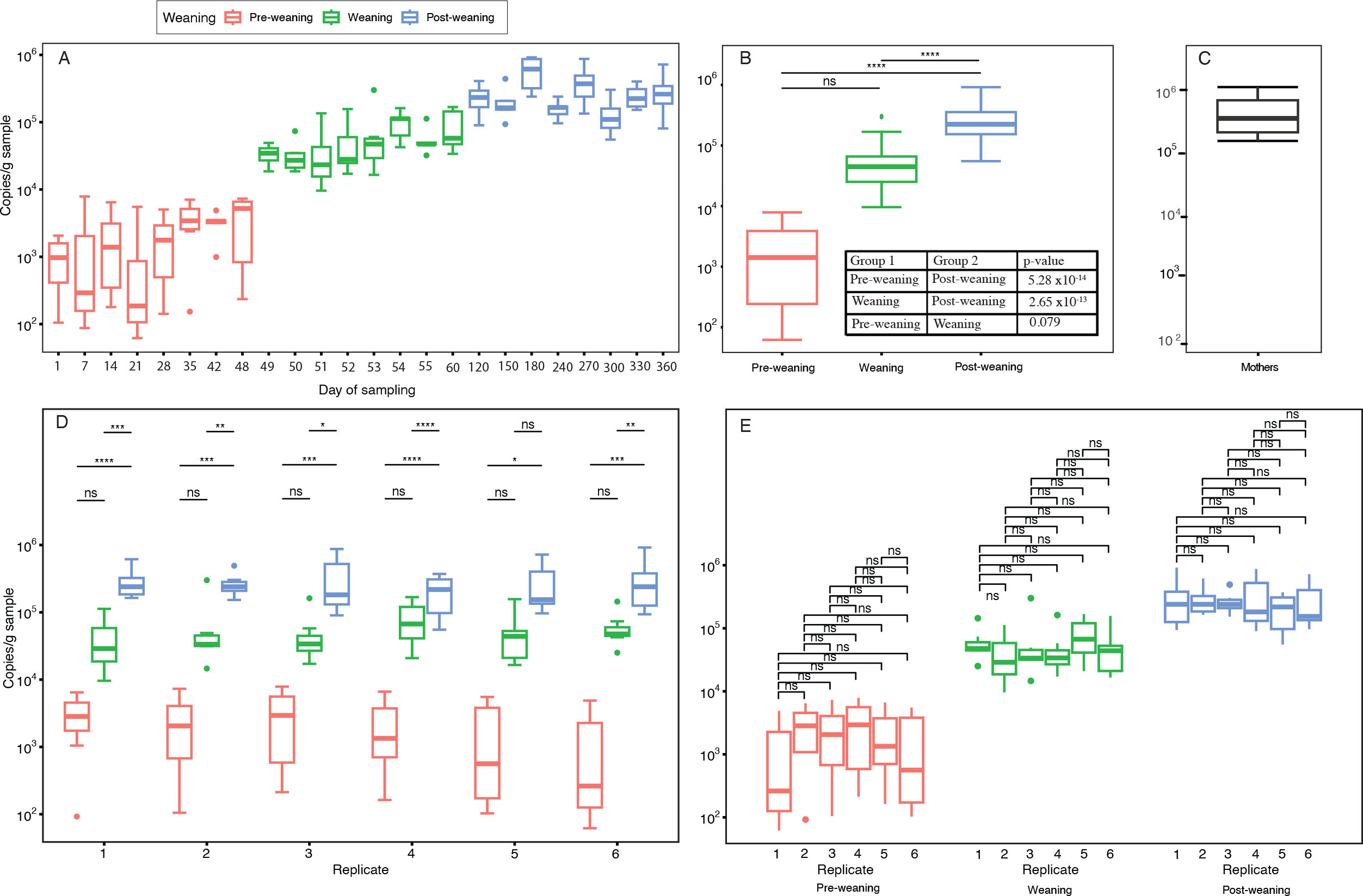
AGF load in the samples studied measured using quantitative PCR. (A) Boxplots showing the progression of the AGF load (expressed as copies/ g feces) as a function of the day of sampling (X-axis). The sampling days are color coded by the weaning phase as shown in the figure legend. The significance of Tukey post-hoc tests for inter-day of sampling comparisons within weaning phases are shown in Table S1. (B) Boxplots showing the progression of the AGF load (expressed as copies/ g feces) as a function of the weaning phase. The significance of Tukey post-hoc test is shown above the box plots and the pairwise comparison p-values are shown in the inset. (C) Boxplot showing the AGF load in mothers (expressed as copies/ g feces). (D) Boxplots showing the AGF load (expressed as copies/ g feces) for each of the calf replicates (X-axis) broken down by the weaning phase. Boxplots are color coded by the weaning phase as shown above (A). The significance of Tukey post-hoc tests for inter-weaning phase comparisons within calves are shown above the box plots, and p-values are shown in Table S2. (E) Boxplots showing the AGF load (expressed as copies/ g feces) for each of the weaning phases (X-axis) broken down by the calf replicate. Boxplots are color coded by the weaning phase as shown above (A). The significance of Tukey post-hoc tests for inter-calf comparisons within weaning phases are shown above the box plots. ns: Not significant.

### Alpha diversity patterns

A total of 928,694 sequences were included in the final analysis. Alpha diversity patterns were assessed using three different indices: Shannon, Simpson, and Inverse Simpson (Figure 2A-F). Analysis of variance showed that the weaning phase significantly affected alpha diversity (p-value <1x10^-13^). Post-hoc Tukey HSD tests for inter- weaning phase comparisons showed that alpha diversity was significantly higher during pre-weaning and weaning phases, when compared to post weaning samples (Figure 2B, D, F) (adjusted p-value < 0.001; Figure 2G). Post-weaning alpha diversity values were also comparable to those in adult cattle (Figure 2B, D, F, G). Within each weaning phase, alpha diversity values did not significantly differ by the day of sampling (Table S3, Figure 2A, C, E), or by calf replicate (Table S4, Figure S1). As well, we calculated the coefficient of variance of alpha diversity measures between different calves sampled at the same timepoint. Pre-weaning and weaning phases were characterized by low levels of inter-calf alpha diversity variability at various time points (Table S5), with a higher level of inter-calf alpha diversity variability observed in calves post-weaning and in adult cattle.

**Figure 2.**
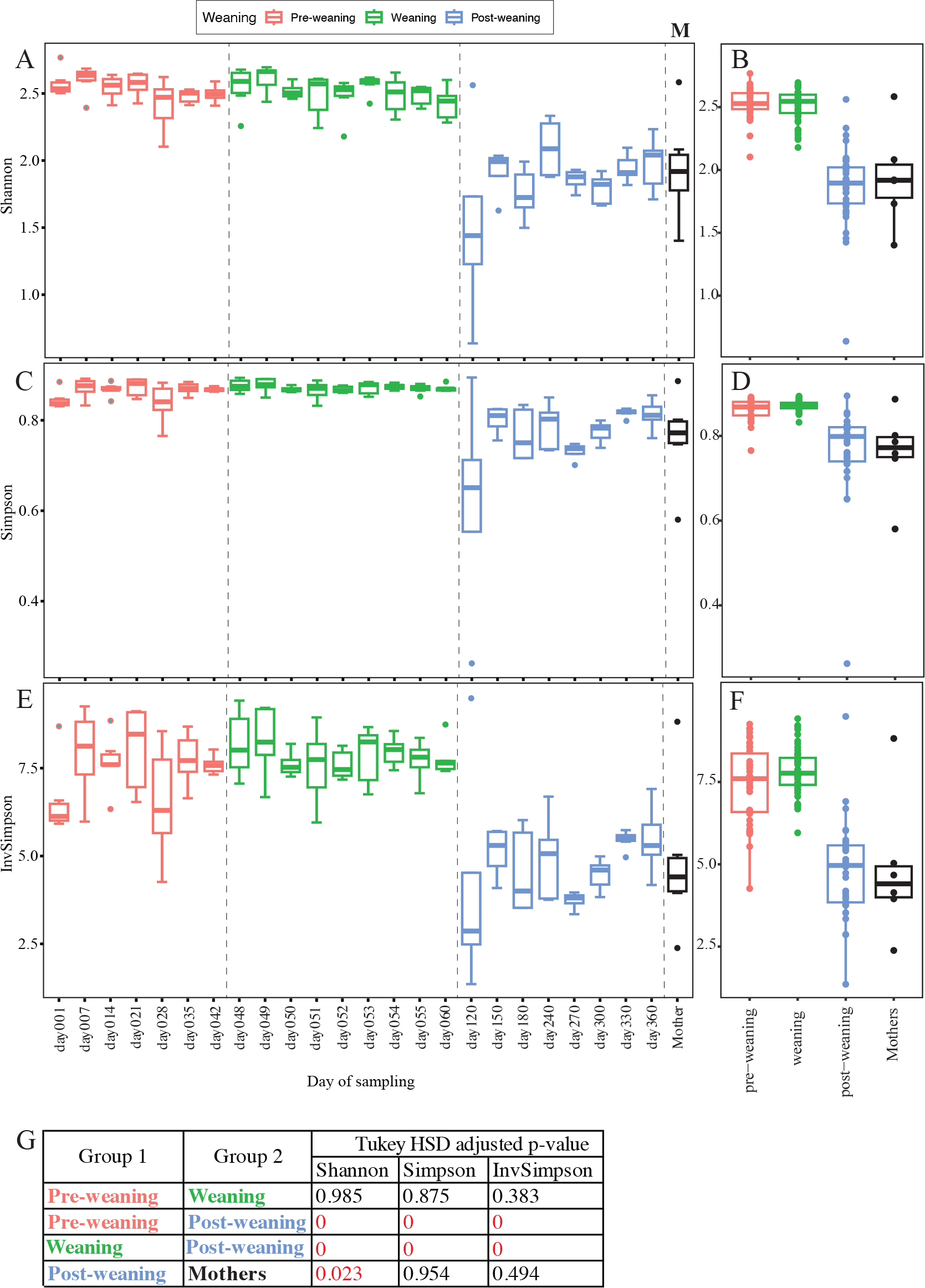
Distribution of alpha diversity measures in the samples studied. Three alpha diversity indices were used: Shannon diversity index (A-B), Simpson evenness index (C-D), and Inverse Simpson (E-F). (A, C, E) Boxplots showing the distribution of Shannon index (A), Simpson index (C), and Inverse Simpson index (E) as a function of the day of sampling (X-axis). The sampling days are color coded by the weaning phase as shown in the figure legend. The significance of Tukey post-hoc tests for inter-day of sampling comparisons within weaning phases are shown in Table S3. (B, D, F) Boxplot showing the distribution of Shannon index (B), Simpson index (D), and Inverse Simpson index (F) as a function of weaning phase. Distribution of alpha diversity indices in the mothers’ samples are shown as the black boxplot in A-F. (G) The significance of Tukey post-hoc tests for inter-weaning phases comparisons.

### Community composition and progression patterns

A total of 84 genera were identified in all datasets examined (Table S6). However, only nine genera (*Caecomyces*, *Neocallimastix*, *Piromyces*, *Orpinomyces*, *Khoyollomyces*, *Cyllamyces*, AL3, NY8, and NY9) were ubiquitous (detected in >50% of samples examined) and abundant (empirically defined as present at >1% relative abundance in >50% of samples examined), with eight more genera ubiquitous (detected in >50% of samples examined) and moderately abundant (empirically defined as present at >1% relative abundance in >10% of samples examined). In addition to these top 17 genera, two genera (*Joblinomyces* and *Feramyces*) exhibited exceptionally high abundance (>5%) in a few samples.

The genus level average percentage abundance (averaged for each sampling date) for these 19 genera is shown in Figure 3A, with individual samples shown in Figure 3B. Discriminable differences in the percentage abundances of specific genera were observed between pre-weaning and weaning phase samples on one hand, and post-weaning samples on the other (Figure 3A, B). Two statistical tests, linear discriminant analysis effect size (LEfSe) and Metastats (Figure 4), were used to detect differentially abundant taxa between pre-weaning (day 1-48), and post-weaning (day 120-360) samples for all 19 genera. Three distinct patterns were observed. Seven genera were significantly more abundant pre-weaning. This group included members of the genera *Khoyollomyces, Orpinomyces*, and the uncultured genera AL3, NY8, NY9, NY6, and NY4 (Figure 4A). Interestingly, some of these genera, e.g. *Khoyollomyces,* AL3, have previously been shown to associate with non-ruminant hindgut fermenters [37]. The most notable decrease in relative abundance during pre-weaning to post-weaning transition was observed for the genus *Khoyollomyces*, with average relative abundance 32.49 ± 5.19% on day 1, decreasing to an average of 0.41 ± 0.02% in the post-weaning phase (*Khoyollomyces* was completely absent from the post-weaning samples at day 180, 300, ad 360). Conversely, six genera were significantly more abundant post-weaning. This group included the genera *Caecomyces*, *Piromyces*, *Cyllamyces*. *Pecoramyces*, *Buwchfawromyces*, and the uncultured genus SK3, many of which are known to be associated with ruminant foregut herbivores [37].

**Figure 3.**
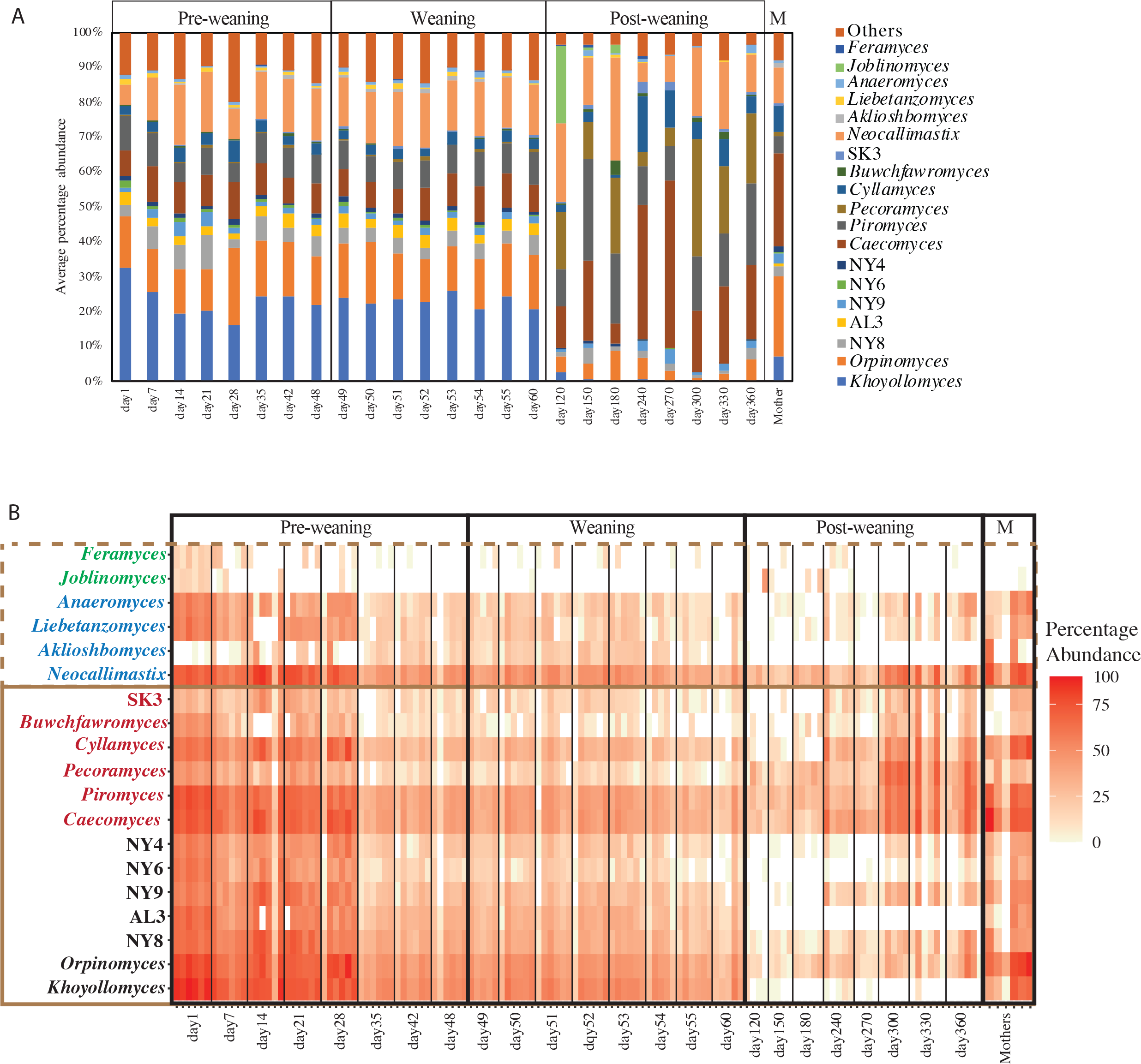
AGF community composition in the samples studied. (A) Average percentage abundance of the 17 ubiquitous (identified in at least 50% of the samples) and abundant (present at >1% relative abundance in at least 10% of samples examined) genera, in addition to the two genera *Joblinomyces* and *Feramyces* that exhibited exceptionally high abundance (>5%) in a few samples. The percentage abundance of these 19 genera was averaged for the six replicate calves for each sampling day (X-axis). The sampling days are grouped by the weaning phase as shown on top of the bar chart. To the right, the average percentage abundance of the 19 genera in the six mothers is shown (labeled “M”). “Others” refer to the collective abundance of all other AGF genera detected in the samples. (B) Heatmap showing the percentage abundance of the same 19 genera in the individual samples with the heatmap key shown to the right. Samples (columns) were grouped first by the sampling day (X-axis) then by weaning phase (thick black borders with labels above the heatmap). To the right, the percentage abundance of the 19 genera in the six mothers is shown (labeled “M” on top). The genera (rows) whose abundances differ significantly by the weaning phase are at the bottom with a solid brown box around and are shown in order starting with genera whose abundances were significantly higher in the pre-weaning phases (black text), then genera whose abundances were significantly higher in the post-weaning phases (red text). The genera (rows) whose abundances did not differ significantly by the weaning phase are on top with a dotted brown box around and include genera with consistent abundance throughout the sampling days (cyan text), and the two genera *Joblinomyces* and *Feramyces* with occasional high abundance in a few samples (green text).

**Figure 4.**
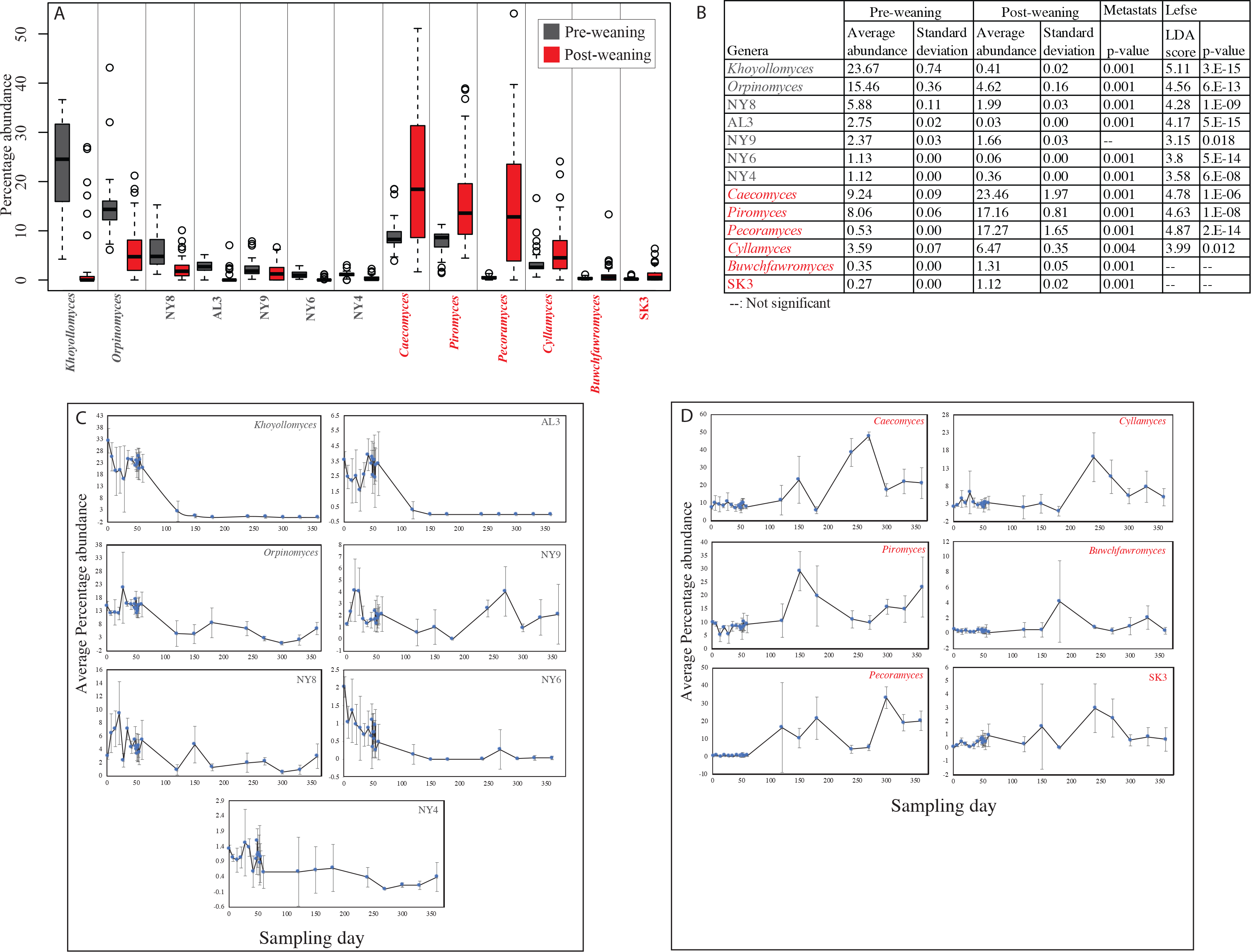
AGF genera with significantly different abundances between pre-weaning and post-weaning phases. (A) Boxplots showing the distribution of the percentage abundances of the 13 genera with significantly higher abundance in pre-weaning (grey boxes), and post-weaning (red boxes) phases. (B) Results of Metastats and linear discriminant analysis (LDA) effect size (LEfSe) analyses to identify genera differentially abundant between the pre-weaning and post-weaning phases. For each genus, the average and standard deviations of abundance is shown for the pre-weaning and post-weaning samples, followed by the methods’ stats, including Metastats p-value, and LEfSe LDA score and p-value. Results are only shown for the genera with at least one significant p-value, and with average percentage abundance >1% in at least one of the two phases. (C-D) Average percentage abundance for each of the thirteen genera whose abundances differ significantly by the weaning phase in the six calves studied as a function of sampling time in days (X-axis). Error bars represent standard deviation. Data are shown for the seven genera that were significantly more abundant during the pre-weaning phase (C), as well as the six genera that were significantly more abundant during the post-weaning phase (D).

The most notable increase in relative abundance during pre-weaning to post-weaning transition was observed for the genera *Percoramyces* (0.53 ± 0.002% pre-weaning to 17.28 ± 1.65% post-weaning), and *Caecomyces* (9.24 ± 0.091% pre-weaning to 23.46 ± 1.97% post-weaning).

For the remaining 6 genera, no statistically significant variation was observed using LEfSe or Metastats. However, two genera showed a sporadic increase in one or more replicates within a single post-weaning sampling time. These included the genus *Joblinomyces* with increased relative abundances in days 120 (one replicate; 85.5%), 150 (one replicate; 1.89%), and 180 (two replicates; 5.33, and 7.87%), and *Feramyces* with increased relative abundances in days 120 (one replicate; 2.27%), 150 (one replicate; 4.26%), and 240 (one replicate; 2.68%) (Figure 3B). The remaining 4 genera showed consistent relative abundance levels regardless of the weaning phase, e.g., the genus *Neocallimastix* with consistently high abundance (>5%), and the genera *Anaeromyces* and *Liebetanzomyces* with consistently moderate to low abundance (0.16-2.5%, and 0.07-1.46%, respectively) (Figure 3A-B).

Comparative analysis between the dams’ AGF community (examined in fecal samples collected on the day of delivery), and calves’ day 1 AGF community (collected within 18h of birth) showed that within individual dam-calf pairs, calves always harbored a higher number of genera compared to their mothers (Day 1 calves harbored 71 to 76 genera, versus 55 to 70 genera in mothers) (Figure 5A-F). However, genera only encountered in calves (i.e., unshared with the dams) always constituted a minor fraction of the community, with only 0.53-1.37% of sequences in the calves’ datasets associated with these genera (Figure 5A-F). More interestingly, when comparing the cohort of dams to the cohort of calves (i.e. pooling all dam samples and all calves samples and comparing both pools), such pattern of higher number of genera in calves was not apparent, with calves collectively harboring 78 genera and mothers harboring 80 genera (Figure 5G).

**Figure 5.**
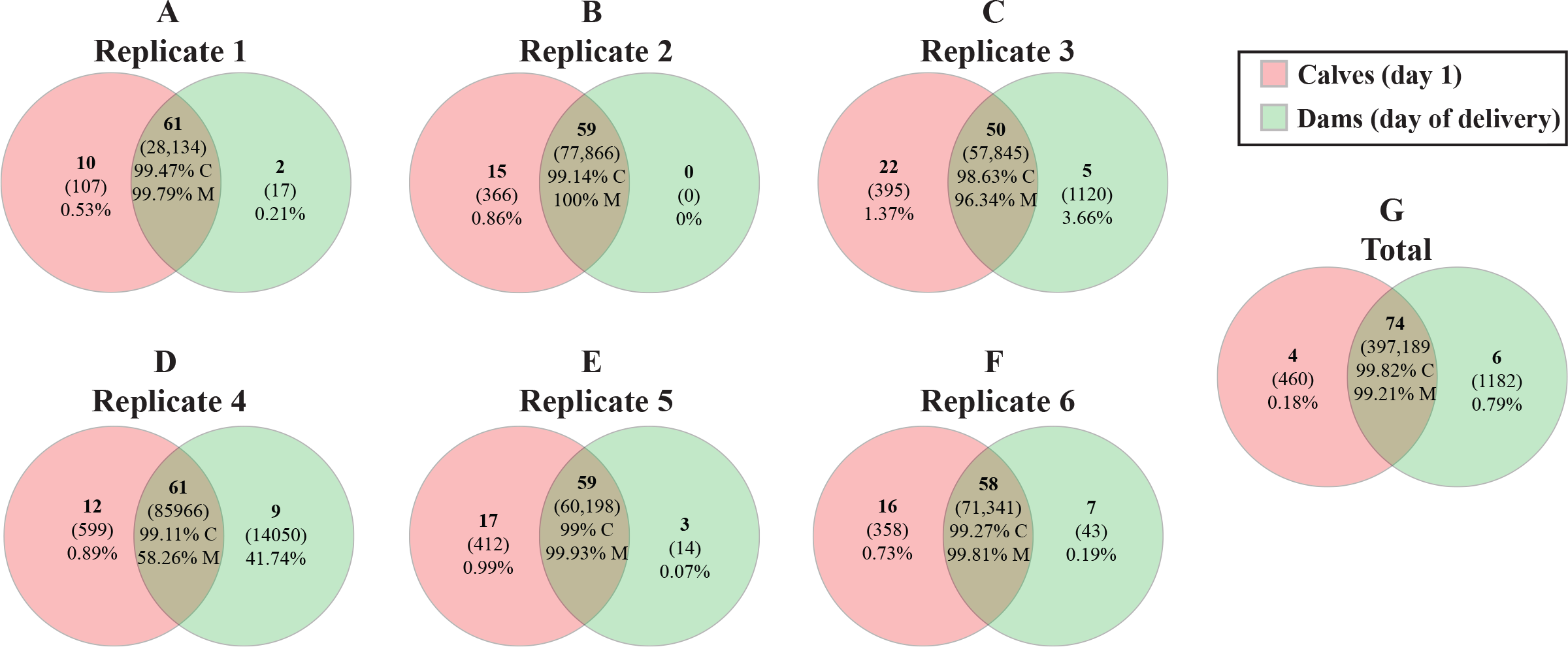
Venn diagrams showing the similarity between the mothers’ AGF community (examined in fecal samples collected on the day of delivery, green), and calves’ day 1 AGF community (pink). (A-F) Venn diagrams for each replicate calf-dam pair are shown. (G) Venn diagram showing the similarity between the cohort of dams to the cohort of calves on day 1 post- birth (i.e. pooling all dam samples and all calves samples and comparing both pools). Shown inside the Venn diagrams are the number of AGF genera (top number in boldface), the number of sequences (the numbers in parentheses), and the percentage of the total community. For the shared communities, two percentages are shown; the percentage of sequences of the total calf community shared between the calf and the dam (percentage followed by “C”), and the percentage of sequences of the total dam community shared between the calf and the dam (percentage followed by “M”).

### Community structure patterns

Patterns of AGF community structure were assessed using principal coordinate analysis plots constructed using the phylogenetic similarity-based index weighted Unifrac (Figure 6). The first two axes explained a high level (72.2%) of the variance observed. Samples clearly clustered by the weaning phase, and within these by the day of sampling (Figure 6A). PERMANOVA analysis (Figure 6B) confirmed that the weaning phase and the sampling day are key predictors of AGF community structure, significantly explaining 53%, and 69% of the variance, respectively. On the other hand, samples did not cluster by the calf replicate (Figure S2), suggesting that temporal progression, but not individual variability, plays a role in shaping the AGF community structure in the studied calves. AGF communities in dams were most closely related to the post-weaning calves’ AGF community (Figure 6A).

**Figure 6.**
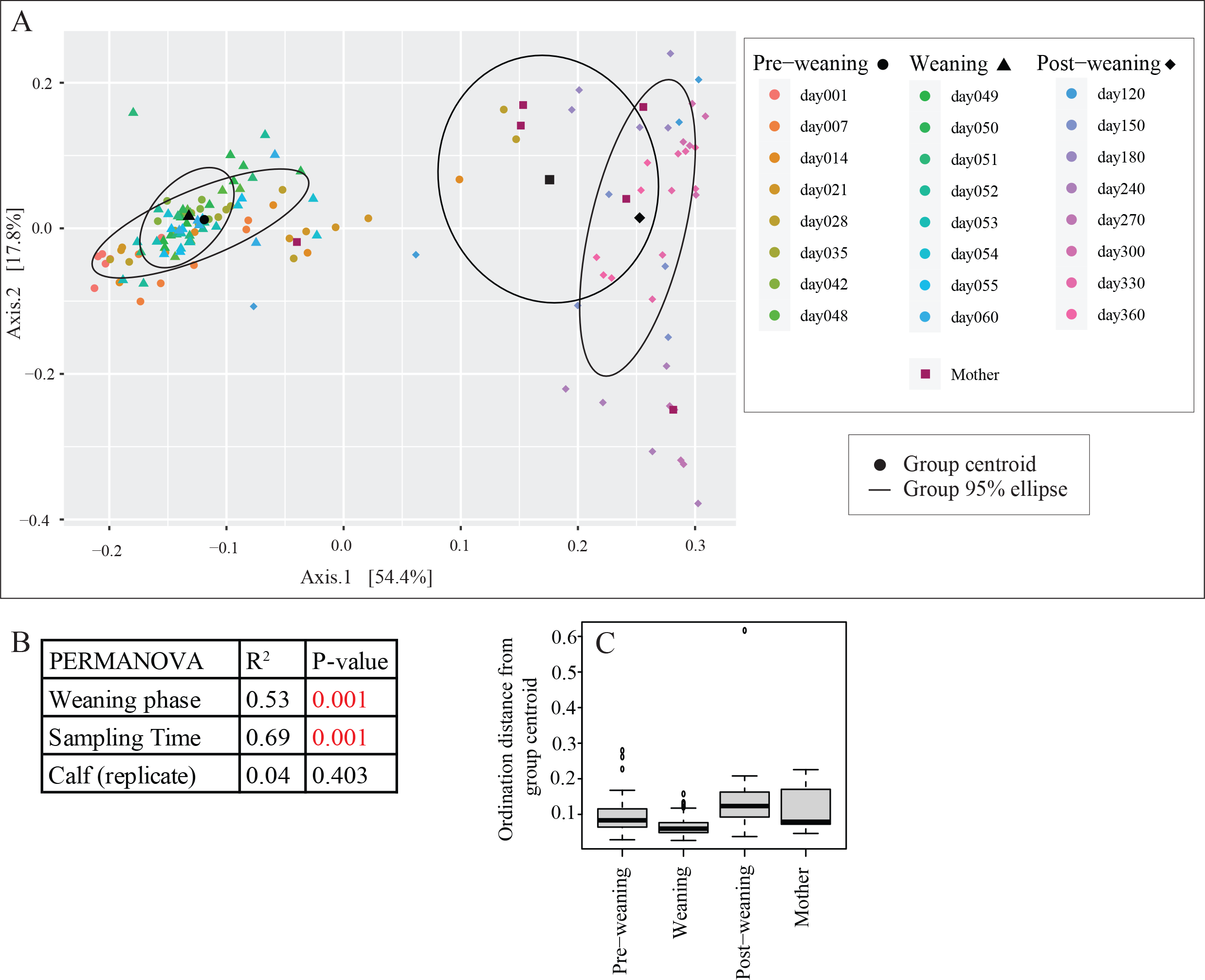
AGF community structure in the samples studied. (A) Principal Coordinate Analysis (PCoA) plot constructed using the pairwise unifrac weighted distance between samples. The first two axes explained 72.2% of the variance. Samples are colored by the day of sampling as shown in the figure legend to the right. Shapes represent the different weaning phases as follows: pre- weaning (•), weaning (..A.), post-weaning (♦), and mothers (▪). Ellipses represent the 95% confidence level for each of the three weaning phases, as well as the dam samples. Each ellipse is centered around the group centroid (black symbols), with slightly larger shapes matching the description above. (B) Results of PERMANOVA tests for partitioning the dissimilarity in community structure among the sources of variation. The PERMANOVA statistic R^2^ depicts the fraction of variance explained by each factor, while the F statistic p-value depicts the significance of each factor in affecting the community structure. (C) Boxplots showing the distribution of each of the samples Euclidean distances on the PCoA shown in (A) from their corresponding group centroids.

Interestingly, comparing the level of variability of different samples within each weaning phase (Figure 6C) demonstrates a higher level of similarity between different samples within pre-weaning and weaning phases (as evidenced by the height of boxplots of the variability of PCoA ordination distance of each sample to its group centroid (Figure 6C)). On the other hand, post-weaning and mother samples were more variable (Figure 6C). Collectively, the results of community composition and community structure analysis demonstrate that the AGF community shifts drastically from a pre-weaning and weaning community enriched in genera commonly associated with non-ruminant hindgut fermenters, to a post-weaning community enriched in genera commonly associated with foregut ruminants.

## Discussion

We followed the AGF community in calves from birth to maturity. Our results provide insights into the mechanism of acquisition, colonization, and selection of individual AGF genera at various GIT developmental stages and associated feeding regimes in cattle. Specifically, we demonstrate that: 1. AGF colonize calves during the first day post-birth at low threshold, and progressively increase in number with calves’ weaning and maturation (Figure 1), 2. AGF communities are significantly more diverse in the pre-weaning and weaning phases compared to post-weaning phases (Figures 2 and S1), 3. AGF community structure post-weaning is drastically different from pre-weaning and weaning communities (Figures 3, 4, and 6), 4. Inter-calf alpha diversity (Figure 2, Table S5) and community structure (Figure 6C) variabilities are lowest in the pre-weaning phases and increase post-weaning, and 5. AGF communities in calves at birth share the majority of their sequences with their mothers (Figure 5), but their communities differ in terms of the relative abundance of different genera (Figures 3 and 4).

We argue that the temporal shifts in AGF load, alpha diversity, and community structure could be understood in the context of calves feeding regiments and GIT development during various stages of maturation. Calves are born as functional non-ruminants, and fed high-protein milk, milk substitute, and protein-rich calf starter feed during pre-weaning. The majority of AGF reside in the rumen in mature cattle (although a minimal presence in cattle hindgut, where 5-10% of microbial carbohydrate degradation is plausible [40]), and specialize in attacking plant biomass in herbivores. As such, we argue that niche and nutritional restrictions severely limit the numbers of AGF during pre-weaning and weaning phases (Figure 1), while the development of a functional niche (developed rumen), as well as the introduction of plant feed (substrate for AGF) post-weaning collectively enrich for the AGF community, resulting in significantly higher numbers in the GIT tract (Figure 1).

Surprisingly, while the AGF load during the pre-weaning and weaning phases (quantified by qPCR) was small, it was unexpectedly diverse (Figure 2), and distinct (Figures 3 and 6) when compared to the post-weaning and adult (dams) communities. Such pattern of higher diversity compared to the dams is consistent with a deterministic negative selection process, where taxa better adapted to the relatively unfavorable conditions in neonatal calves immediately post birth (i.e., absence of functional rumen, lack of plant biomass, and relatively higher oxygen levels) are selected from the dam’s community. This negative selection process, occurring during the first 24 hours of birth, results in the suppression of specific highly abundant genera typically active in the rumen, hence simultaneously decreasing the AGF load (Figure 1), while increasing community evenness and overall alpha diversity (Figure 2). Further, the smaller and more diverse community selected at this stage bears little resemblance to the dams and the post-weaning community (Figure 6), and, surprisingly, is characterized by higher relative abundance of some genera typically associated with hindgut fermenters (Figure 4), e.g. genera *Khoyollomyces* (the most abundant AGF genus in pre-weaning and weaning samples), and AL3, both typically encountered in the GIT tract of the hindgut fermenter family Equidae [20, 37, 41]. Such pattern suggests that calves’ hindgut might provide a haven for AGF in the calves’ early life stages, prior to the development of a functional rumen. Fermentation processes are known to occur in the hindgut of cattle, even post-weaning [40].

Weaning and transition to solid plant diet resulted in a decrease in diversity (Figure 2), and a clear shift in AGF community composition (Figure 3) and structure (Figure 6). These changes are indicative of a positive selection process, where the development of a functional rumen, as well as the introduction of abundant plant biomass substrates selectively enrich for specific genera more adapted to fast growth in the rumen, hence decreasing evenness and overall alpha diversity. The majority of genera selected (*Caecomyces*, *Piromyces*, *Pecoramyces*, *Cyllamyces*, *Buwchfawromyces*) have previously been shown to occur in higher relative abundance in foregut ruminants [37]. Further the evolution of these genera has recently been tied to the evolution of a functional rumen in their hosts [37]. However, the specific differences in physiological preferences, metabolic capabilities, and genomic content underpinning selective preference of various AGF genera to specific gut types and/or developmental stage are currently unclear.

The current study followed AGF progression in six different calves from day 1 after birth to maturity. In addition to providing adequate replication for statistical analysis, this setting allowed the quantification of inter-calf variabilities in AGF load, diversity, and community structure across various stages of calves’ development. Surprisingly, during pre-weaning phases, lower levels of inter-calf variability were observed in AGF load (Figure 1, coefficient of variance ranging between 3.52-9.17%), as well as in alpha (Figures 2, S1, Table S4), and beta diversity (Figure 6D), when compared to post-weaning samples. A possible explanation for such an unexpected pattern is that negative selection in calves during pre-weaning phases uniformly inhibits growth of specific genera due to specific factors (e.g. oxygen sensitivity) that do not vary between calves replicates; resulting in a more uniform community. On the other hand, positive selection based on competition for resources during post-weaning phases could be influenced by various stochastic processes (e.g. priority arrival), or that slight inter-calf variabilities (e.g. occasional illness, slight variability in feed intake) could influence inter-AGF competition and community dynamics and trajectories. This could explain the sporadic increase in the abundance of a few genera in one or more replicates within a single post-weaning sampling time, e.g. *Feramyces* and *Joblinomyces* (Figure 3B)

Sampling mother fecal samples on the delivery day provide valuable insights onto dam- calf transmission and AGF community establishment post-birth. AGF are clearly confined to GIT in calves [11], and hence other sources of transmission, e.g. placenta, umbilical cord, colostrum, and amniotic fluid, as previously suggested for bacteria [3], appear highly unlikely. Our results indicate that the dam of a specific calf could provide the majority of AGF taxa for neonatal colonization (Figure 5A-F), with possible additional input from other dams and animals present within the same location (Figure 5G) Finally, it is worth noting that the observed patterns of AGF progression differ from those observed with bacterial and archaeal communities in key aspects. A relatively large body of knowledge exists on bacterial temporal progression in cattle, and few common themes have emerged from these studies [3, 42–45]. All studies suggest a rapid increase and establishment of a high load of microbial community in the calves in the first 24 hours, in contrast to the persisting low numbers in AGF observed here. In addition, bacterial communities during pre-weaning phases are less diverse than those post-weaning, while the reverse was observed with AGF in this study. Finally, bacterial communities post-weaning are more stable, with much lower variations between replicates compared to pre-weaning phases, again in contrast to the pattern observed here with AGF [46]. These differences could mainly be rationalized in light of the great collective metabolic and physiological versatility of the unicellular bacteria and archaea when compared to AGF, allowing rapid growth, colonization, and propagation during all phases of growth and in all locations within the calves’ GIT tract. The rapid colonization post-birth [47] is a reflection of the fact that the presence of oxygen in the GIT of neonatal calves, as well as the lack of rumen, and dependence on milk (or milk replacer) and high protein feed starter during pre-weaning phases are by no means an impediment to bacterial growth, but rather select for microorganisms adapted to such substrates, niches, and conditions as previously demonstrated [43]. This is in contrast to the fact the AGF growth is impeded under such conditions. The progressive increase in bacterial diversity could best be understood by the complexity and high specialization of bacterial members in food webs operating to degrade complex substrates under anaerobic conditions. Specifically, the transition to plant biomass necessitates the development of a complex community of bacteria, with highly specialized members mediating degradation of complex plant polysaccharides (e.g. cellulose, hemicellulose, pectin), fermentation of oligomers and sugar monomers into volatile fatty acids (VFAs), recycling cell detritus, syntrophic metabolism of intermediate VFAs, and hydrogen and CO_2_ conversion to acetate and methane, among others [48]. This is in contrast to AGF specialization in the initial attack on plant biomass and plant polymer degradation [11].

In conclusion, our results suggest a pattern of rapid AGF establishment in calves GIT post-birth, driven by early negative selection of an AGF community adapted for survival in the calves’ undeveloped GIT tract and liquid high-protein diet intake. The transition to plant feed and the associated development of rumen as the main organ for fermentation post-weaning induces a positive selection process, resulting in a marked increase in AGF load, decrease in diversity, and a shift to AGF genera adapted to rumen. Further studies on specific differences in physiological preferences and metabolic capabilities of various AGF taxa are needed to understand the basis for preferences of various genera to specific weaning phases. Such knowledge, together with the baseline understanding of AGF community progression outlined here, could open the way towards rational manipulation of AGF communities towards better health and growth outcomes in cattle.

## Contributions

Conceptualization and experimental design by NHY and MSE. Sample collection, archiving, and laboratory experimentation ALJ, JC, EP, DM, and APF. Data analysis by ALJ, JC, EP, APF, and NHY. Manuscript writing by ALJ, APF, MSE, and NHY.

