## Supplementary document for "Temporal progression of anaerobic fungal communities in dairy calves from birth to maturity"

**Health protocol for calves**

Upon arrival at the OSU dairy, the calves were orally administered a modified-live rota-coronavirus vaccine (Calf-Guard; Zoetis Animal Health, Parsipanny, NJ, USA). On day 7, all calves were banded for castration and administered a vaccination for *Clostridium perfringens* types C and D and *Clostridium tetani* (Bar-Vac CD/T; Boehringer Ingelheim Animal Health USA Inc., Duluth, GA, USA). Calves were revaccinated on day 152. On day 11, all calves were given a direct-fed microbial bolus (Tri-Start Jr.; Accelerated Genetics, Baraboo, WI, USA).

On day 11, two calves developed scours and were treated with 1.89 L of electrolytes (Re-Sorb; Zoetis Animal Health) and milk replacer was reduced by half through day 19. On day 15, the two calves developed a fever and were treated by a veterinarian and one calf received intravenous antibiotics (Nuflor, Merck & Co., Inc., Rahway, NJ, USA) and the other received Nuflor subcutaneously. The calf receiving intravenous Nuflor was administered a second dose on day 19. On day 13 and 14, all calves were given electrolytes and milk replacer was reduced by half and were administered a probiotic oral gel (Probios Bovine One Oral Gel, Vet Plus Inc., Menomonie, WI, USA) on day 17. Two calves developed scours on day 28 and were given electrolytes once daily through day 32.

On day 74, calves were treated with Amprolium (Corid 9.6% oral solution; Huvepharma Inc., Peachtree City, GA, USA) drench due to a nearby calf not associated with this study developing coccidiosis. On day 112, all calves received topical ivermectin (Ivomec pour-on, Boehringer Ingelheim Animal Health USA Inc.) as part of normal husbandry practices. Calves received Bovishield Gold FP5 (Zoetis Animal Health) and Vision 7 (Merck & Co., Inc.) on day 159. Additionally, on day 159, four of the calves were dehorned by a veterinarian and received meloxicam (Unichem Pharmaceuticals USA, East Brunswick, NJ, USA).

**Supplementary Figures:**

**Figure S1.** Distribution of alpha diversity measures within the calves. Boxplots showing the distribution of Shannon index (A), Simpson index (B), and Inverse Simpson index (C) as a function of the weaning phase (X-axis). Boxplots are color coded by calf as shown in the figure legend. Distribution of alpha diversity indices in the mothers’ samples are shown as the black boxplot.

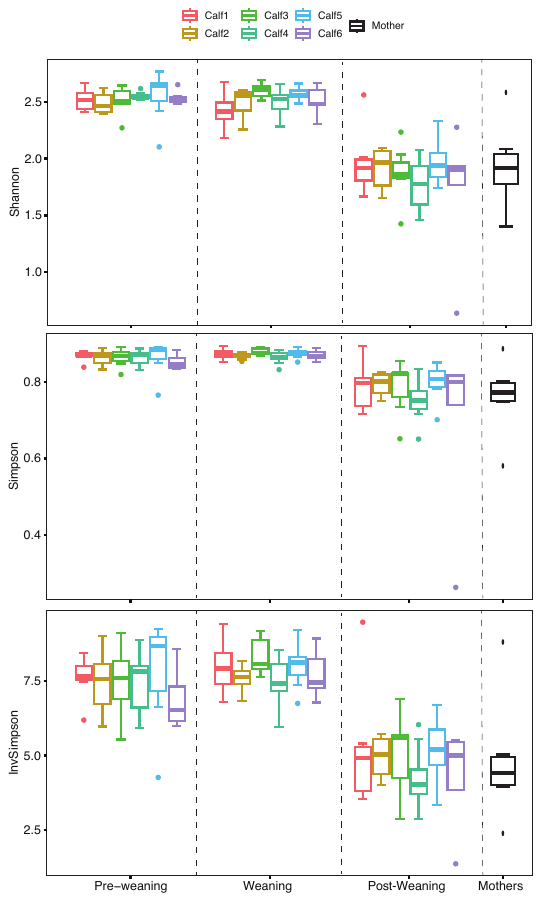

**Figure S2.** Principal Coordinate Analysis (PCoA) plot constructed using the pairwise Unifrac weighted distance between samples. The first two axes explained 72.2% of the variance. Samples are colored by the calf replicate as shown in the figure legend to the right. Shapes represent the different weaning phases as follows: pre-weaning (), weaning (), post-weaning (), and mothers (). Samples clearly clustered by the weaning phase, and within these by the day of sampling as shown in Figure 6A, but not by the calf replicate.

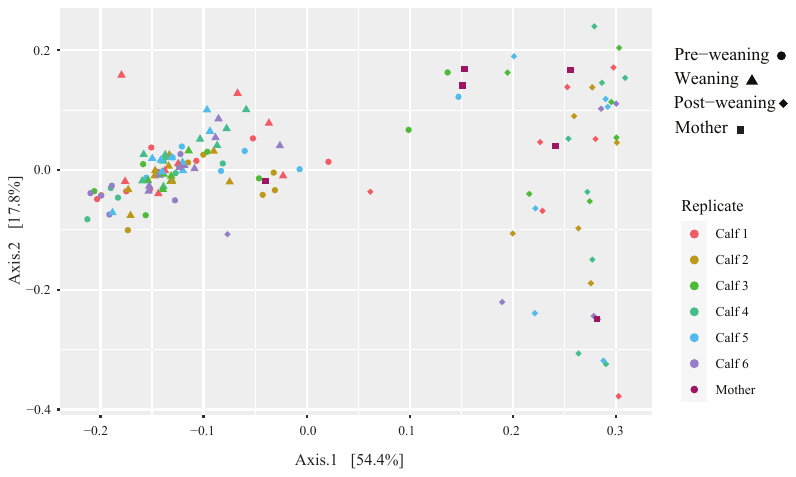

Supplementary tables.

**Table S1.** The significance of Tukey post-hoc tests for comparison of AGF copies/g feces between different day of sampling within each of the weaning phases.

**Table S2.** The significance of Tukey post-hoc tests for comparison of AGF copies/g feces between different weaning phases within each of the calves replicates studied.

**Table S3.** The significance of Tukey post-hoc tests for comparison of alpha diversity measures between different day of sampling within each of the weaning phases. Distribution of alpha diversity results are shown in Figure 2A, C, and E, for Shannon, Simpson, and Inverse Simpson indices, respectively.

**Table S4.** The significance of Tukey post-hoc tests for comparison of alpha diversity measures between calf replicates within each of the weaning phases. Distribution of alpha diversity results are shown in Figure S1.

**Table S5.** Average, standard deviation, and coefficient of variation for alpha diversity measures between different calves sampled at the same timepoint (grouped by the weaning phases).

**Table S6.** Community composition in the samples studied. Samples are grouped by the day of sampling and results are shown for each of the calf replicates, as well as the mothers on the day of delivery. Heatmap colors correspond to the percentage abundance and are scaled from red (0%) to green (highest abundance).
